# Hydroxychloroquine-mediated inhibition of SARS-CoV-2 entry is attenuated by TMPRSS2

**DOI:** 10.1101/2020.07.22.216150

**Authors:** Tianling Ou, Huihui Mou, Lizhou Zhang, Amrita Ojha, Hyeryun Choe, Michael Farzan

## Abstract

Hydroxychloroquine, used to treat malaria and some autoimmune disorders, potently inhibits viral infection of SARS coronavirus (SARS-CoV-1) and SARS-CoV-2 in cell-culture studies. However, human clinical trials of hydroxychloroquine failed to establish its usefulness as treatment for COVID-19. This compound is known to interfere with endosomal acidification necessary to the proteolytic activity of cathepsins. Following receptor binding and endocytosis, cathepsin L can cleave the SARS-CoV-1 and SARS-CoV-2 spike (S) proteins, thereby activating membrane fusion for cell entry. The plasma membrane-associated protease TMPRSS2 can similarly cleave these S proteins and activate viral entry at the cell surface. Here we show that the SARS-CoV-2 entry process is more dependent than that of SARS-CoV-1 on TMPRSS2 expression. This difference can be reversed when the furin-cleavage site of the SARS-CoV-2 S protein is ablated. We also show that hydroxychloroquine efficiently blocks viral entry mediated by cathepsin L, but not by TMPRSS2, and that a combination of hydroxychloroquine and a clinically-tested TMPRSS2 inhibitor prevents SARS-CoV-2 infection more potently than either drug alone. These studies identify functional differences between SARS-CoV-1 and -2 entry processes, and provide a mechanistic explanation for the limited *in vivo* utility of hydroxychloroquine as a treatment for COVID-19.

**Author Summary:** The novel pathogenic coronavirus SARS-CoV-2 causes COVID-19 and remains a threat to global public health. Chloroquine and hydroxychloroquine have been shown to prevent viral infection in cell-culture systems, but human clinical trials did not observe a significant improvement in COVID-19 patients treated with these compounds. Here we show that hydroxychloroquine interferes with only one of two somewhat redundant pathways by which the SARS-CoV-2 spike (S) protein is activated to mediate infection. The first pathway is dependent on the endosomal protease cathepsin L and sensitive to hydroxychloroquine, whereas the second pathway is dependent on TMPRSS2, which is unaffected by this compound. We further show that SARS-CoV-2 is more reliant than SARS coronavirus (SARS-CoV-1) on the TMPRSS2 pathway, and that this difference is due to a furin cleavage site present in the SARS-CoV-2 S protein. Finally, we show that combinations of hydroxychloroquine and a clinically tested TMPRSS2 inhibitor work together to effectively inhibit SARS-CoV-2 entry. Thus TMPRSS2 expression on physiologically relevant SARS-CoV-2 target cells may bypass the antiviral activities of hydroxychloroquine, and explain its lack of *in vivo* efficacy.

## Introduction

The pandemic coronavirus disease 2019 (COVID-19, caused by SARS coronavirus 2 (SARS-CoV-2) poses serious threat to global public health [1]. In the first months following the onset of the pandemic, several existing drugs were recommended to repurposed for treatment of COVID-19, among them chloroquine and its derivative hydroxychloroquine sulfate (hydroxychloroquine) [2]. The use of hydroxychloroquine has become controversial as clinical trials suggest that this drug is ineffective as either a treatment or a prophylaxis against SARS-CoV-2 infection. The United States Food and Drug has since revoked its emergency use authorization for this drug [3–7]. This disappointing result contrasts with its promising cell culture studies which demonstrated a half-maximal effective concentration (EC_90_) of 6.9 μM against SARS-CoV-2 replicating in Vero E6 cells. This concentration is achievable *in vivo* with a well-tolerated 500 mg daily administration and similar with the EC_50_ of the relatively more successful remdesivir (1.76 μM) [8–10].

Hydroxychloroquine has been suggested to restrict multiple steps in the coronaviral lifecycle [11–13], but its inhibitory effect on viral entry as a lysosomotropic agent is best defined [14]. It elevates endosomal pH and subsequently interrupts activities of cathepsin L, one of the entry factors for coronaviruses [15, 16]. Coronaviral entry requires both receptor engagement and fusion activation by proteolytic processing the S glycoprotein. The SARS-CoV-1 and −2 S proteins bind angiotensin-converting enzyme 2 (ACE2), their common receptor [17–19]. Two obligate proteolysis sites for fusion activation have been identified within the S proteins, namely at the junction of the S1 and S2 domain, and at the S2’ site in an exposed loop of the S2 domain [20]. The SARS-CoV-2 S1/S2 junction is cleaved in virus producing cells by proprotein convertases that cleave a distinctive furin-recognition site at this boundary [21]. In contrast, the SARS-CoV-1 S1/S2 boundary is cleaved in the virus target cell after receptor engagement by either cell-surface TMPRSS2 or endosomal cathepsin L [15, 22, 23]. The S2’ sites of both viruses are cleaved in the target cell, again by either TMPRSS2 or cathepsin L (**Fig. 1**). These proteolysis events and ACE2-binding prime the S protein for conformational changes that mediate fusion of the viral and cellular membranes [22–25]. While hydroxychloroquine is known to suppress cathepsin proteolysis activity, its impact on TMPRSS2-mediated viral entry is unknown.

**Figure 1.**
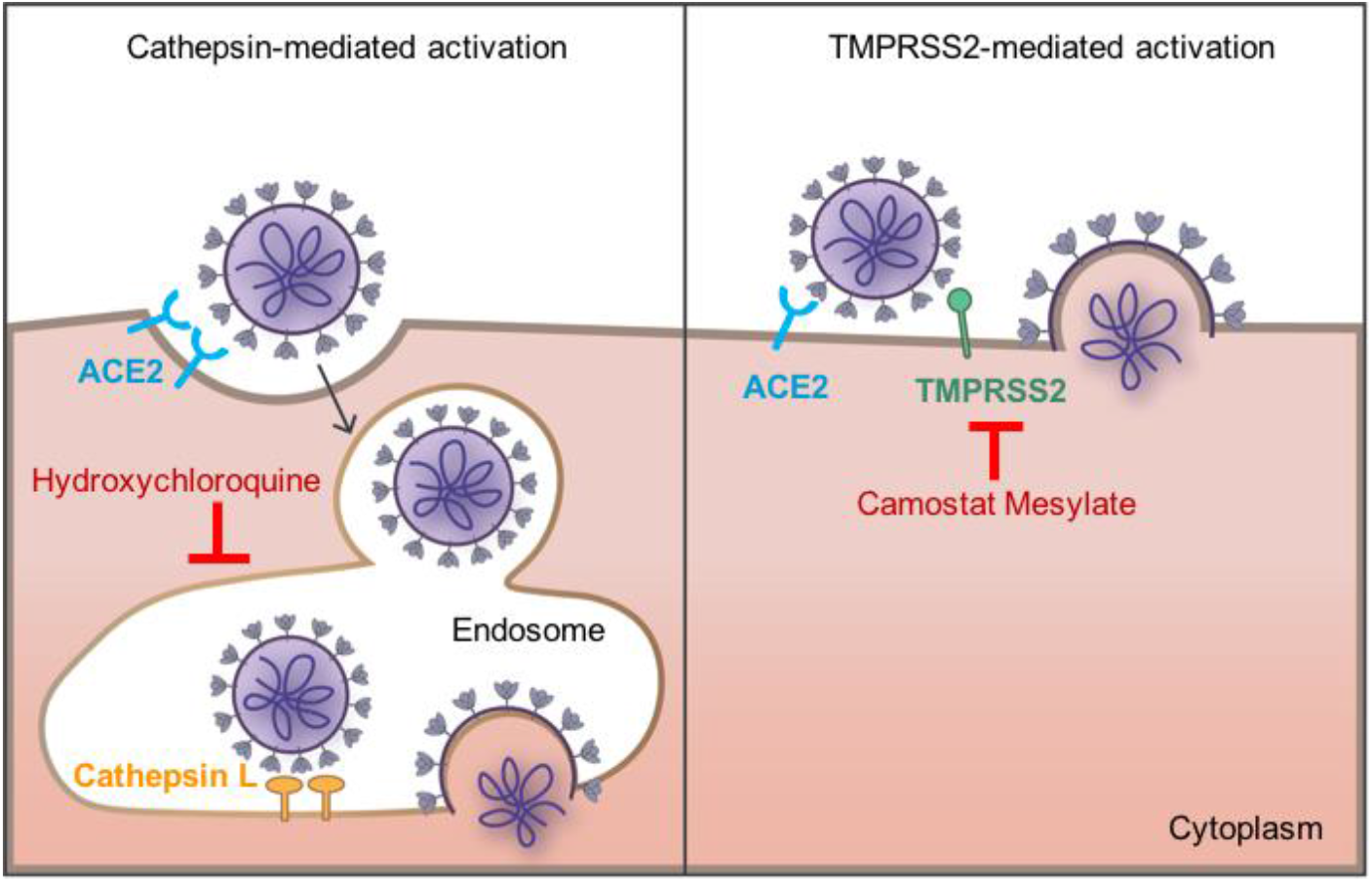
SARS-CoV-1 and SARS-CoV-2 fusion can be activated by either or both of two pathways. The coronaviruses bind the cellular receptor ACE2 and then must be activated by proteolysis with either a surface-expressed protease like TMPRSS2 or by cathepsin L in the endosome. Only cathepsin L-mediated proteolysis requires endosomal acidification. Camostat mesylate inhibits TMPRSS2 activity, whereas hydroxychloroquine, like ammonium chloride, inhibits endosomal acidification.

Cells that express both ACE2 and TMPRSS2 are present in multiple tissues including lung (alveolar and bronchial), buccal mucosa, nasal mucosa, ileum, colon, and myocardium epithelium [23, 26]. Recent studies of single-cell RNA sequencing further locate these highly susceptible cells in respiratory tree, cornea, esophagus, ileum, colon, gallbladder and common bile duct [27]. Three major cell types are identified to have TMPRSS2 and ACE2 co-expression: lung type II pneumocytes, ileal absorptive enterocytes, and nasal goblet secretory cells [28]. Of note, while ACE2 has a generally lower expression level and a narrower distribution than TMPRSS2, most ACE2-positive cells in the respiratory tract also express TMPRSS2 [27–31].

Here we evaluated the infectivity of SARS-CoV-1 and -2 S proteins on cells in the presence and absence of TMPRSS2. We show that TMPRSS2 expression has a markedly greater impact on entry mediated by the SARS-CoV-2 S protein than by that of SARS-CoV-1. We further show that antiviral efficiency of hydroxychloroquine on SARS-CoV-2 S-protein-mediated entry is negatively impacted by the expression level of TMPRSS2. Consistent with these observations, when cells are expressing a high level of TMPRSS2, the TMPRSS2 inhibitor camostat was more potent than hydroxychloroquine at inhibiting SARS-CoV-2 infection, but the converse for SARS-CoV-1. Finally, we demonstrate that the anti-SARS-CoV-2 activity of hydroxychloroquine could be enhanced by camostat but not by compounds that inhibit cathepsin L. Thus failure of hydroxychloroquine in clinical studies reflects the presence of TMPRSS2 in key tissues and its importance to the SARS-CoV-2 entry process.

## Results

### TMPRSS2 allows efficient cell entry mediated by the SARS-CoV-2-S protein, bypassing a cathepsin L-dependent endosomal entry pathway

TMPRSS2 can proteolytically activate membrane fusion of a variety of respiratory viruses including influenza A virus, SARS-CoV-1, MERS-CoV and SARS-CoV-2 [22–24, 32, 33]. TMPRSS2 expression was shown to have an impact on the susceptibility of cells to MERS-CoV [32]. We first investigated how TMPRSS2 expression affected SARS-CoV-2 infectivity. To do so, we measured the viral entry of pseudovirions (PV) bearing SARS-CoV-1 or SARS-CoV-2 S proteins (SARS1-PV, SARS2-PV). HEK293T-ACE2 cells (a stable cell line) transiently transfected with TMPRSS2 or control plasmids (mock) were used as target cells. As demonstrated in **Fig 2A**, when TMPRSS2 was expressed, viral entry mediated by the SARS-CoV-2-S protein was significantly increased (up to 1000-fold). Indeed, the same amount of infection was observed when one-sixteenth of the SARS2-PV was used in the presence of TMPRSS2. In contrast, enhancement of SARS1-PV entry by TMPRSS2 was less pronounced (0.5-10 fold). We then asked whether TMPRSS2 expression allowed SARS-CoV-2-S protein-mediated entry to bypass the endosomal activation pathway. To test this possibility, TMPRSS2(−) and TMPRSS2(+) cells were treated with ammonium chloride prior to infection. As expected, the inhibitory effect of ammonium chloride was more robust on TMPRSS2(−) cells for both SARS1- and SARS2-PV (**Fig 2B**). Without TMPRSS2, more than 60% of the SARS2-PV entry was inhibited by ammonium chloride. For VSV-G pseudotypes, which are pH sensitive but do not utilize TMPRSS2 for proteolysis activation, the inhibition efficiency of ammonium chloride was moderate, and unaffected by TMPRSS2 expression. TMPRSS2-expressing cells treated by ammonium chloride has a 10-fold higher viral infection compared to the control group (TMPRSS2(−) cells with DMSO treatment), suggesting that for SARS-CoV-2 S-protein mediated entry, the TMPRSS-mediated activation pathway is much more efficient than the endosomal activation pathway. Moreover, in the presence of TMPRSS, ammonium chloride had only a modest effect on SARS2-PV infection.

**Figure 2.**
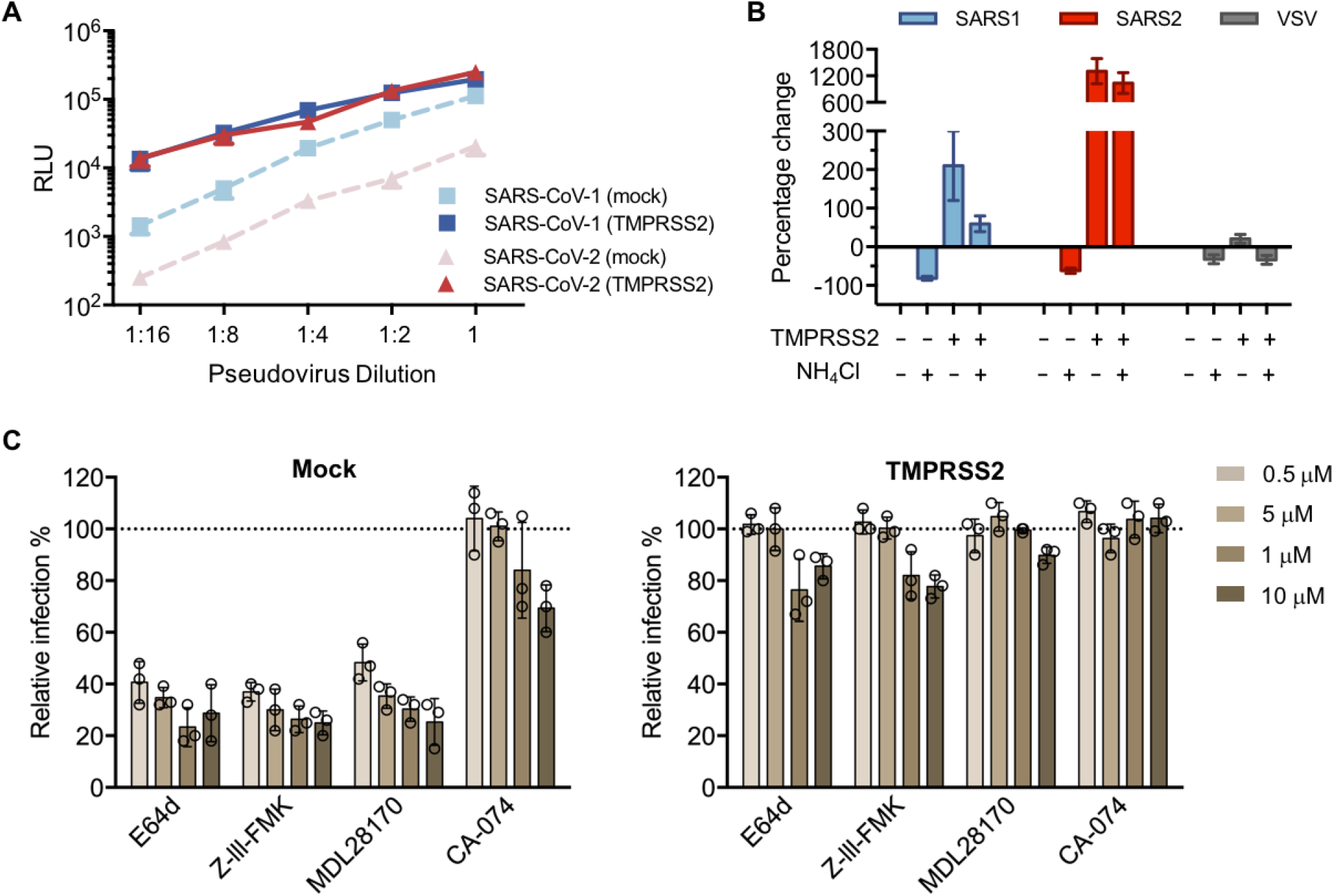
Suppressing endosomal proteases activities does not reduce SARS-CoV-2 infectivity on TMPRSS2-expressing cells. **(A)** 293T-ACE2 cells were transfected with a vector control plasmid (TMPRSS2-negative), or a TMPRSS2 expression plasmid, 24 hours prior to infection. Luciferase reporter retroviruses pseudotyped with SARS-CoV-1 (SARS1), SARS-CoV-2 (SARS2) spike proteins were used for infection at titers yielding equivalent infection in the presence of TMPRSS2. Pseudovirus was serially diluted by 2-fold starting from stock (RLU≈250,000 for TMPRSS2-expressing cells). Cells were inoculated with diluted pseudovirus, and viral entry was determined by the luciferase activity in cell lysates within 48 hours post infection. Note that luciferase expression from SARS-CoV-2 pseudotypes on TMPRSS2-negative cells (RLU≈20,000) could be achieved by approximately 16-fold dilution of the same amount of virus on TMPRSS2-expressing cells. **(B)** 293T-ACE2 cells were transfected with a vector control plasmid, or a TMPRSS2 expression plasmid. Retroviruses pseudotyped with SARS-CoV-2, SARS-CoV-1 spike proteins, and VSV-G were used for infection. Cells were treated by 50 mM of ammonium chloride before infection. Infection of TMPRSS2-negative cells without treatment was used for normalization. Bars indicate the means (±SD) of three independent experiments. **(C)** A panel of cathepsin inhibitors was tested on 293T-ACE2 cells transiently expressing a control plasmid or TMPRSS2 prior to infection. E64d and Z-III-FMK inhibit both cathepsin B and cathepsin L; MDL28170 specifically inhibits cathepsin L, and CA-074 inhibits cathepsin B. Cells were treated with the indicated concentrations of protease inhibitors or DMSO for 2 hours, then inoculated with luciferase retrovirus harboring SARS-2-CoV spike proteins. Luciferase activity was measured at 48 hours post inoculation. Relative infection (%) was calculated from infection of DMSO-treated cells.

To determine the precise proteases involved in the SARS-CoV-2-S proteolysis activation, a panel of cysteine protease inhibitors were tested on TMPRSS2(−) and TMPRSS2(+) HEK293T-ACE2 cells. Among these compounds, E64d and Z-III-FMK inhibit both cathepsin B and cathepsin L; MDL281740 is cathepsin L inhibitor and CA-074 is cathepsin B inhibitor. In the absence of TMRPPS2 expression, E64d, Z-III-FMK, and MLD281740 showed robust inhibition of SARS2-PV (**Fig 2C**). These results suggest that, as observed with SARS1-PV, cathepsin L but not cathepsin B, facilitatea SARS-CoV-2 infection. In contrast, when TMPRSS2 was expressed, the antiviral activity of these cathepsin inhibitors was largely abrogated. Collectively, these results suggest that in the absence of TMPRSS2 expression, endosomal cathepsin L is critical to viral entry mediated by SARS-CoV-2-S protein; however, when TMPRSS2 is expressed, the role of cathepsin L is markedly diminished. They further indicate that TMPRSS2 contributes more strongly to SARS-CoV-2 infection than to SARS-CoV-1.

### TMPRSS2 expression significantly attenuates the antiviral effect of hydroxychloroquine against SARS-CoV-2-S

Hydroxychloroquine can be used as a broad-spectrum antiviral drug against multiple viruses [34]. Although its exact antiviral mechanism remains unclear, it is well established that hydroxychloroquine accumulates within the acidic organelles such as endosomes [14]. As a weak base, it thereby increases the pH of endosomes, and subsequently interferes with the activities of pH-dependent endosomal protease. Because the antiviral effect of ammonium chloride on SARS-CoV and SARS-CoV-2 can be overcome by TMPRSS2 expression, we investigated whether TMPRSS2 expression also affects the antiviral activities of hydroxychloroquine. We observed that transient expression of TMPRSS2 in HEK293T-ACE2 cells resulted in higher TMPRSS levels than stable expression (data not shown), although only a low plasmid concentration was used for transient transfection (5 ng/well in 96-well plate). To demonstrate the effect of different amount of TMPRSS2 on antiviral activities of hydroxychloroquine, both transient (high) and stable (low) TMPRSS2 HEK293T-ACE2 cells were used as targets. For HEK293T-ACE2 cells transiently expressing the control plasmids (**Fig 3A**), hydroxychloroquine potently inhibited viral entry of SARS1- and SARS2-PV (IC_50_=1.53 μM and 1.62 μM, respectively). These IC_50_ values are consistent with a previous cell culture study, using replicative SARS-CoV-2 on Vero cells, which do not express TMPRSS2 [10]. However, when TMPRSS2 was expressed, hydroxychloroquine-mediated inhibition of SARS2-PV was substantially attenuated. The IC_50_ of hydroxychloroquine against pseudotyped SARS-CoV-2 was increased by 5- to 60-fold for low and high TMPRSS2-expressing cells, respectively (**Fig 3B**). SARS1-PV also became more resistant to hydroxychloroquine with TMPRSS2 expression. Its IC_50_ was increased by 20-fold with high TMPRSS2 expression, although no difference was observed with low TMPRSS2 expression. Hydroxychloroquine modestly reduced VSV-G-mediated cell entry, and TMPRSS2 expression did not affect this inhibition.

**Figure 3.**
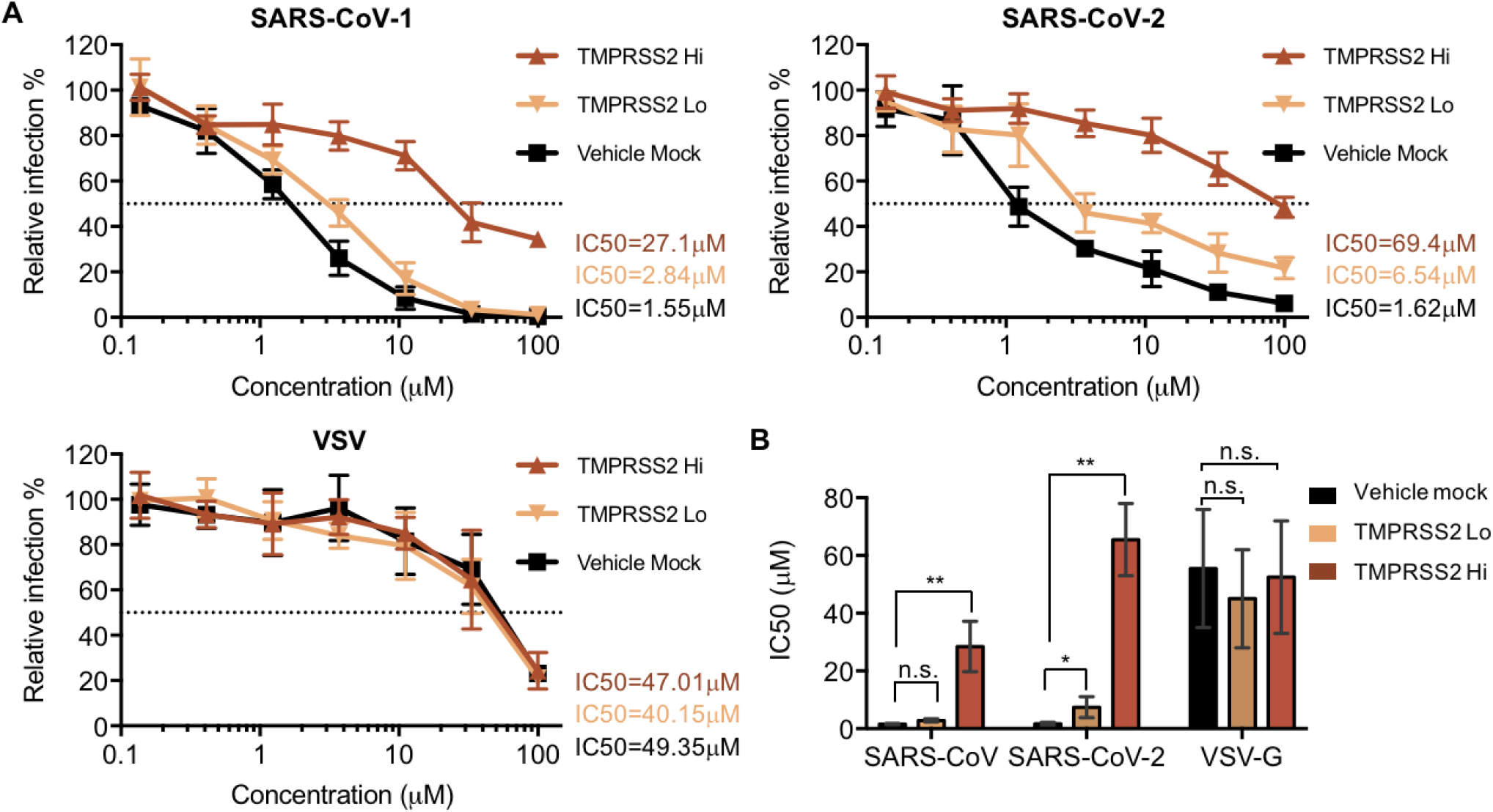
Antiviral effect of hydroxychloroquine is dependent on TMPRSS2 expression. **(A)** A stable cell line was generated from 293T-ACE2 cells to express low amount of TMPRSS2 (TMPRSS2 Lo, black solid line). 293T-ACE2 cells were transiently transfected with a vector control (vehicle mock, grey solid line), or TMPRSS2 (TMPRSS2 Hi, red solid line). The cells were treated with hydroxychloroquine or DMSO before virus inoculation. The results are presented as a percentage of infection of DMSO-treated cells (≈ 250,000 RLU for all three pseudotypes from TMPRSS-expressing cells.) **(B)** A summary of IC_50_ changes of pseudoviruses under different conditions. Data shown represent the means (±SD) of three independent experiments. Unpaired Student’s t-test was used to assess the statistical significance of the difference between IC_50_ on mock transfected cells (TMPRSS2-negative) and TMPRSS2-positive cells. (**: P < 0.01. *: P < 0.05. n.s.: P > 0.05.)

### Suppression of TMPRSS2 restores the antiviral efficiency of hydroxychloroquine

To confirm that the antiviral effect of hydroxychloroquine on SARS-CoV-1 and SARS-CoV-2 is masked by TMPRSS2 activities, we next explored whether suppression of TMPRSS2 could rescue the antiviral efficiency of hydroxychloroquine. Camostat, a clinically proven drug that specifically inhibits TMPRSS2, was tested in combination with hydroxychloroquine on TMPRSS2-expressing cells incubated with SARS1-, SARS2-, or VSV-G-PV (**Fig 4A**). Notably, hydroxychloroquine alone inhibited SARS1-PV than camostat, whereas the reverse was true for SARS-CoV-2, where camostat alone inhibited much more efficiently than hydroxychloroquine. The combination of both drugs inhibited SARS1- and SARS2-PV more effectively than either drug alone. We also examined the effect of varying the concentration of hydroxycholoroquine in the presence of fixed (10 μM) amounts of camostat, the cathepsin-L and cysteine protease inhibitor E64D, or DMSO alone (**Fig 4B**). Consistent with the presumption that E64D and hydroxycholoquine redundantly interferes with endosomal activation of the S protein, E64D only modestly altered the inhibitory activity of hydroxychloroquine for SARS2-PV (changing the hydroxychloroquine IC_50_ from 69.4 μM to 25.0 μM) and SARS1-PV (27.1 μM to 9.4 μM). In contrast, the same fixed amount of camostat had a more than 20-fold impact on hydroxychloroquine inhibition of SARS2-PV (changing the IC_50_ from 69.4 μM to 3.2 μM), but much less impact on SARS1-PV (27.1 μM to 10.6 μM). These data again demonstrate that SARS2-PV are more responsive to TMPRSS2 inhibition than are SARS1-PV, and that TMPRSS2 must be inhibited for hydroxychloroquine to be effective.

**Figure 4.**
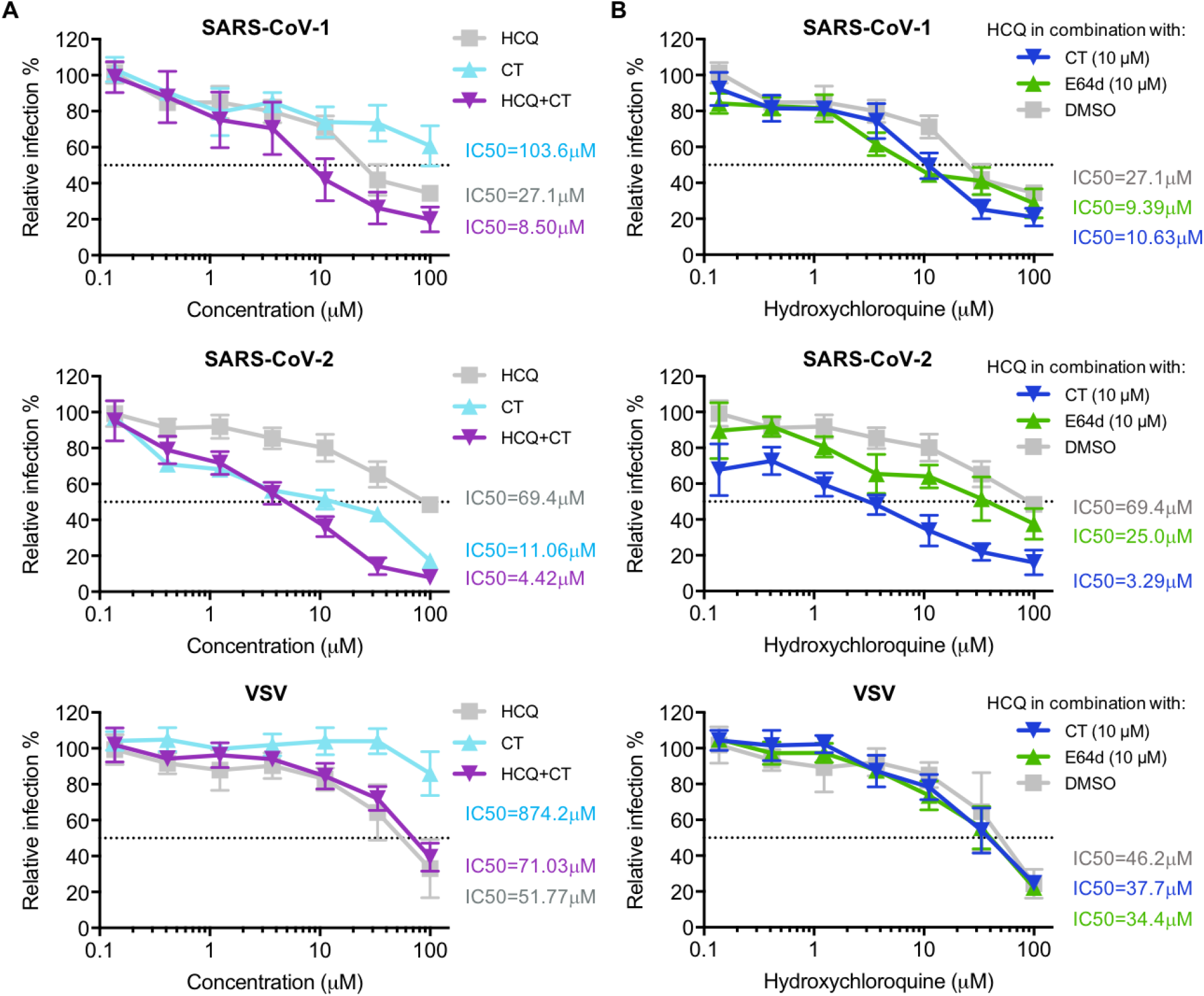
Suppression of TMPRSS2 restores the antiviral efficiency of hydroxychloroquine. **(A)** Hydroxychloroquine (HCQ) and camostat (CT) were tested on 293T-ACE2 cells transiently expressing TMPRSS2 prior to infection. The antiviral efficiency of each drug alone and the combination of two drugs were compared. Cells were challenged with retrovirus pseudotyped with SARS-CoV-2, SARS-CoV-1 spike proteins, and VSV-G after drug or DMSO treatment. Luciferase activity was measured at 48 hours post inoculation. Relative infection (%) was calculated from infection of DMSO-treated cells. **(B)** Cells were treated with drugs or DMSO prior to pseudovirus inoculation. HCQ was serially diluted in complete media (grey solid line), complete media containing 10 μM of camostat (CT, blue solid line), or 10 μM of E64d (cyan solid line), respectively. Relative infection (%) was calculated from infection of DMSO-treated cells.

### An S-protein furin-cleavage site increases reliance on TMPRSS2 expression

The data from experiments above consistently indicated that SARS-CoV-2 is more dependent on TMPRSS2 than SARS-CoV-1. Because the SARS-CoV-2 S protein differs from that of SARS-CoV-1 by the presence of a furin-cleavage site at its S1/S2 boundary, we investigated whether ablating the furin site (FKO) decreases its reliance on TMPRSS2 expression. In addition, previous studies suggest that the furin site renders SARS-CoV-2 S protein relatively unstable, a phenotype partially restored by a naturally occurring D614G mutation in the S1 domain. We therefore investigated whether this D614 mutation would render it less dependent on TMPRSS2. The infectivity pseudoviruses bearing the FKO and D614G S-protein variants was compared to wildtype SARS-CoV-2 S protein on TMPRSS2-positive and TMPRSS2-negative cells. As shown in **Fig 5A**, when pseudovirus titers were adjusted to be equivalent in the absence of TMPRSS2 (left panel), entry mediated by the wild-type S protein was most enhanced by TMPRSS2 expression (up to 100-fold), and that mediated by the furin knock-out FKO was least affected (10-fold). The D614G variant showed an intermediate phenotype (30-fold), consistent with its greater stability than wild-type S protein. To confirm this results, we then adjusted pseudovirus titers to be equivalent in the presence of TMPRSS2 (right panel). Consistent with the previous results, entry mediated by wild-type S protein was most affected by the absence of TMPRSS2, the FKO mutant was least affected, and the D614G variant again exhibited an intermediate phenotype. The drug sensitivity of these two S-protein mutants to hydroxychloroquine and camostat was also compared to the wild-type S protein on both TMPRSS2-negative and TMPRSS2-positive cells (**Fig 5B**). In the absence of TMPRSS2, entry mediated by all three S-protein variants was similarly sensitive to hydroxychloroquine, and, as expected, unaffected by camostat. However, in the presence of TMPRSS2, the FKO S protein was more inhibited by hydroxychloroquine and less inhibited by camostat, suggesting that this variant relies more on S-protein activation in the endosome than do both naturally occurring S proteins bearing furin sites. Thus the SARS-CoV-2 S-protein furin-cleavage site determines its greater reliance on TMPRSS2, its greater sensitivity to camostat, and its greater resistance to hydroxychloroquine.

**Figure 5.**
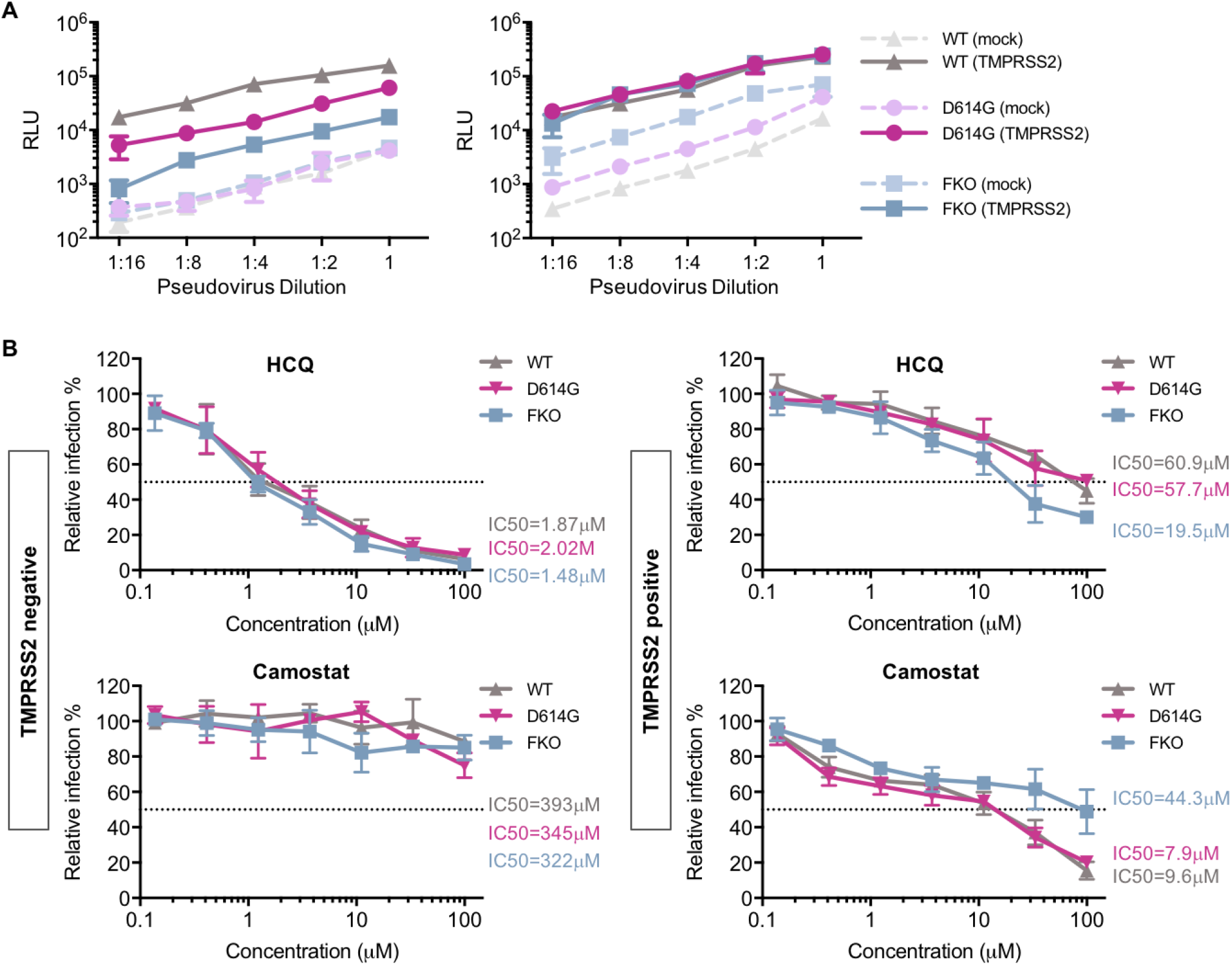
Furin cleavage in the virion producer cell correlates with TMPRSS2 dependence. **(A)** An experiment similar to that in Fig 1A except that viral infectivity of pseudoviruses bearing the SARS-CoV-2 wildtype (WT) S protein, a D614G S-protein variant, or furin-site knockout (FKO) S protein was determined on 293T-ACE2 cells with or without transient expression of TMPRSS2. Pseudovirus titers were adjusted so that they were equivalent in the absence (left panel) or presence (right panel) of TMPRSS2. **(B)** The effect of TMPRSS2 on the antiviral efficiency of hydroxychloroquine (HCQ) and camostat was compared. 293-ACE2 cells were challenged with retrovirus pseudotyped against wildtype, D614G mutation, and FKO SARS-CoV-2 pseudoviruses after drug or DMSO treatment. Luciferase activity was measured at 48 hours post inoculation. Relative infection (%) was calculated from infection of DMSO-treated cells.

## Discussion

It has been previously demonstrated that SARS-CoV-1 and SARS-CoV-2 entry into cells depends on the expression on two somewhat redundant proteases, namely cathepsin L located in acidic cellular compartments, and TMPRSS2, expressed on the plasma membrane [24, 35]. It is further understood from these and other observations that, unlike pH-dependent viruses such as influenza A virus, conformational changes of the spike protein of coronaviruses are not dependent on pH [15]. Rather, proteolytic activation by cathepsin L is dependent on endosomal acidification, and thus elevating endosomal pH prevents cathepsin-L-mediated entry. However, the TMPRSS2-mediated entry pathway, in which S-protein is presumed to be activated at the plasma membrane, is not affected by pH (**Fig 1**). Here we extend these observations in two directions. First we show that SARS-CoV-2 is more dependent on TMPRSS2 than is SARS-CoV-1, and that this difference can largely be explained by the presence of a furin cleavage site in the SARS-CoV-2 S protein. Second, we show that TMPRSS2 expression overcomes the antiviral effect of hydroxychloroquine, thus providing a mechanistic explanation for its poor therapeutic efficacy against SARS-CoV-2 despite encouraging cell-culture results.

We present multiple lines of evidence that SARS-CoV-2 is more sensitive to the presence of TMPRSS2 than is SARS-CoV-1. With pseudovirus (PV) titers adjusted so that SARS1-PV and SARS2-PV infection were comparable in the presence of TMPRSS2, SARS2-PV transduced cells markedly less efficiently in its absence (**Fig 2A**). In the presence of TMPRSS2, SARS1-PV are more sensitive than SARS2-PV to inhibitors of endosomal acidification such as ammonium chloride (**Fig 2B**) and hydroxychloroquine (**Fig 3A and 3B**). Specifically, the IC_50_ of hydroxychloroquine for SARS1-PV was three-fold lower than for SARS2-PV with high TMPRSS2 expression, while their sensitivities to hydroxychloroquine are equivalent in the absence of TMPRSS2 (**Fig 3A and Fig 4A**). By contrast, SARS2-PV were 12-fold more sensitive to the TMPRSS2 inhibitor camostat (**Fig 4A**). Although TMPRSS2-mediated pathway is preferred over the cathepsin-L-mediated pathway for both SARS-CoV-1 and −2, these data indicate that SARS-CoV-1 utilizes the cathepsin-L-pathway more efficiently than SARS-CoV-2, whereas SARS-CoV-2 is more dependent on TMPRSS2 than SARS-CoV-1.

What accounts for this difference? The most obvious difference between the SARS-CoV-1 and -2 S proteins is the presence of a polybasic or furin-cleavage site at the boundary between the S1 and S2 domains [21]. This furin site is present in other human coronaviruses, for example MERS-CoV and HCoV-OC43 [36, 37], but it has not previously been observed in any of the many SARS-like coronaviruses identified in bats [38]. Thus, the S1/S2 boundary of the SARS-CoV-2 S protein, but not that of SARS-CoV-1, is cleaved at this site in the virus-producing cells. Indeed, when the SARS-CoV-2 furin-cleavage site is ablated, the mutant PV is less impacted by the addition or removal of TMPRSS2 compared to the wildtype (**Fig 5A**), and these pseudoviruses are relatively more sensitive to hydroxychloroquine-mediated inhibition, and less sensitive to camostat (**Fig 5B**).

Thus furin-cleavage in the virus-producing cell correlates with greater dependence on TMPRSS2, and lower dependence on cathepsin L. If one assumes that several proximal S-protein trimers must be fully cleaved and activated to mediate fusion, this difference is relatively easy to understand. Specifically, furin cleavage in virus-producing cells reduces the number of required proteolytic events in the target cell, and thus in many cases TMPRSS2 alone can fully activate fusion. In the case of SARS-CoV-1, or when the furin-cleavage site is replaced by one cleaved only by TMPRSS2, more target-cell cleavage events are required. In such cases, to complete the proteolytic activation of sufficient numbers of S proteins, cathepsin L-mediated proteolysis, and therefore viral endocytosis and endosomal acidification, are necessary.

However our data also make clear that, in the absence of TMPRSS2, SARS-CoV-2 utilizes cathepsin L less efficiently than SARS-CoV-1. For example, when titers are adjusted so that infections by SARS1- and SARS2-PV are identical in the presence of TMPRSS2 (**Fig 1A**), their efficiencies are markedly different in its absence. This difference can perhaps be explained by the relative instability of the wild-type SARS-CoV-2 S protein [16, 39] in two ways. First, the furin-cleaved SARS-CoV S protein has been shown to prematurely shed [40], resulting in fewer S proteins per virion and pseudovirion. The resulting lower density of S proteins per virion may impair the ability of cross-linked ACE2 to promote endocytosis of the virus. Alternatively, the less stable S protein may be further destabilized in the acidifying endosome, disrupting the ordered steps of fusion mediated by proteolytic activation and conformational transitions of S2. Thus the more stable D614G SARS-CoV-2 S protein variant [40] is modestly less affected by the presence and absence of TMPRSS2 than the wild-type (D614) S protein (**Fig 5A**).

The greater dependence of SARS-CoV-2 on TMPRSS2 has an immediate implication for the treatment of COVID-19. Specifically, it implies that inhibitors of endosomal acidification will have less impact on SARS-CoV-2 in the presence of TMPRSS2. We show here that indeed, TMPRSS2 helps bypass the hydroxychloroquine-mediated inhibition of SARS2-PV infection. Most physiologically relevant target cells in the body, include type II pneumocytes and ciliated nasal epithelial cells express TMPRSS2. Thus the potent inhibition of SARS-CoV-2 by hydroxychloroquine in Vero E6 cells, where TMPRSS2 is largely absent, overestimated its potency by 10- to 40-fold, depending on TMPRSS2 expression (**Fig 3A**). However, our results suggest that some efficacy of hydroxychloroquine could be restored if TMPRSS2 is inhibited by camostat (**Fig 4B**), an observation directly relevant to clinical trials, for example NCT04338906, combining the two inhibitors.

In summary, we show that the inhibitory effect of hydroxychloroquine on SARS-CoV-2 entry is attenuated by TMPRSS2, thus explaining its limited clinical efficacy. We further show that SARS-CoV-2 is more dependent on TMPRSS2 and less dependent on cathepsin L than SARS-CoV-1, and that these differences are likely due to the presence of a furin-cleavage site in the S protein of SARS-CoV-2. Finally we show inhibition of both TMPRSS2 and cathepsin L may be necessary to fully block virus entry in cells that express both proteases.

## Materials and methods

### Plasmid

Plasmids encoding TMPRSS2 and the control empty vectors were purchased from OriGene (SC323858, PS100020). DNA sequence of SARS-CoV-2 S protein (GenBank YP_009724390) was codon-optimized and synthesized by Integrated DNA Technologies (IDT, Coralville, IA, USA), and was cloned subsequently into pCAGGS vector using In-Fusion® HD Cloning Kit (Takara Bio USA) according to manufacturer’s instructions. Furin-site mutated SARS2-S was created by replacing -RRAR- at S1/S2 junction with -SRAS-; the SARS-CoV D614G and furin-site mutated S proteins were made by site-directed mutagenesis.

### Pseudovirus production

MLVs pseudotyped with variant spike or envelope proteins were generated as described before [41]. Briefly, HEK293T cells were co-transfected with three plasmids, pMLV-gag-pol, pQC-Fluc and pCAGGS-SARS2-S-cflag or pcDNA3.1-SARS1-S or pCAGGS-VSV-G, and the medium was refreshed after 6h incubation of transfection mix. The supernatant with produced virus was harvested 72h post transfection and clarified by passing through 0.45μm filter. Clarified viral stocks were supplemented with HEPES with the final concentration of 10mM and stored at −80°C for long-term storage.

### Cell culture and stable cell line

The HEK293T cell line expressing human ACE2 (hACE2) were created by transduction with produced vesicular stomatitis virus (VSV) G protein-pseudotyped murine leukemia viruses (MLV) containing pQCXIP-myc-hACE2-c9 as described before. Briefly, HEK293T cells were co-transfected with three plasmids, pMLV-gag-pol, pCAGGS-VSV-G and pQCXIP-myc-hACE2-c9, and the medium was refreshed after overnight incubation of transfection mix. The supernatant with produced virus was harvested 72h post transfection and clarified by passing through 0.45μm filter. The parental 293T cells were transduced with generated MLV virus, and the 293T-hACE2 cell lines were selected and maintained with medium containing puromycin (Sigma). hACE2 expression was confirmed by immunofluorescence staining using mouse monoclonal antibody against c-Myc antibody 9E10 (Thermo Fisher) and Goat-anti-mouse FITC (Jackson ImmunoResearch Laboratories, Inc).

HEK293T/ACE2/TMPRSS2 stable cell line was also constructed by transducing 293T-hACE2 cell line with MLV pseudovirus made by cotransfection of pMLV-gag-pol, pQCXIB-TMPRSS2-Flag and pCAGGS-VSV-G at 3:2:1 ratio into 293T cells. Stable cell lines were maintained in Dulbecco’s Modified Eagle Medium (DMEM, Life Technologies) at 37°C in a 5% CO_2_-humidified incubator. Growth medium were supplemented with 2 mM Glutamax-I^™^ (Gibco), 100 μM non-essential amino acids (Gibco), 100 U/mL penicillin and 100 μg/mL streptomycin (Gibco), and 10% FBS (Gibco). For the 293T-ACE2 stable cell line, 3 μg/mL of puromycin was added to the growth medium to maintain expression of ACE2. For 293T/ACE2/TMPRSS2, 1 μg/mL of puromycin and 10 μg/mL blasticidin was added to the growth medium. To measure surface TMPRSS2 expression, cells were detached by 1mM EDTA in PBS and then stained by 4 ug/ml of anti-Flag M2 antibody (Sigma, F1804) and 2 ug/ml of Goat anti-mouse IgG (H+L) conjugated with Alexa 647 (Jackson ImmunoResearch Laboratories, Inc, Cat# 115-606-146).

### Pseudovirus infection

HEK293T-ACE2 cells were seeded at 30% density in poly-lysine pre-coated 96-well plates 12-15 hours prior to transfection. Cells in each well were then transfected with 0.3 μL of lipofectamine 2000 (Life Technologies) in complex with 5 ng of a vector control plasmid or a plasmid encoding TMPRSS2. Cell culture medium was refreshed at 6 hours post transfection. Additional 18 hours later, cells in each well were infected with pseudovirus diluted in 100 μL of culture medium containing 2% FBS (Gibco). Cells were spin-infected at 4°C for 30 min at 3000xg to allow virus-binding to cells, followed by 2 hours of incubation at 37°C. After incubation with virus, supernatant was removed and each well was replenished with 200 μL of fresh media containing 2% FBS.

### Luciferase assay for viral entry

At 48 hours post infection, cells were lysed in wells and subjected to *Firefly* luciferase assays. Viral entry was determined using Britelite Plus (PerkinElmer), and luciferase expression was measured using a Victor X3 plate reader (PerkinElmer). For experiments testing drug inhibition, target cells were treated by the chemicals diluted in 100 μL of media containing 2% FBS to the final indicated concentrations. After incubation for 2 hours at 37°C, supernatant was removed before virus transduction.

### Statistical analysis

Data expressed as mean values ± S.D. or S.E.M, and all statistical analysis was performed in GraphPad Prism 7.0 software. IC_50_ of drugs was analyzed using default settings for log(inhibitor) vs. normalized response method. Statistical difference was determined using non-paired Student’s t-test. Differences were considered significant at P < 0.05.

## Author contributions

T.O., H.M., L.Z., A.O., H.C., and M.F. designed experiments and analyzed the results. T.O., H.M., L.Z., A.O. execute the experiments. T.O. and M.F. conceived the study and wrote the manuscript.

